# Integrating genome-wide association and transcriptome predicted model identify novel target genes with osteoporosis

**DOI:** 10.1101/771543

**Authors:** Peng Yin, Muchun Zhu, Fan Hu, Jiaxin Jiang, Li Yin, Shuqiang Wang, Yingxiang Li

## Abstract

Osteoporosis (OP) is a highly polygenetic disease which is usually characterized by low bone mineral density. Genome-wide association studies (GWAS) have identified hundreds of genetic loci associated with bone mineral density. However, the biological mechanisms of these loci remain elusive. To identify potential causal genes of the associated loci, we detected trait-gene expression associations by transcriptome-wide association study (TWAS) method. It directly imputes gene expression effects from GWAS data, using a statistical prediction model trained on GTEx reference transcriptome data, with blood and skeletal tissues data. Then we performed a colocalization analysis to evaluate the posterior probability of biological patterns: association characterized by a single shared causal variant or two distinct causal variants. The ultimate analysis identified 276 candidate genes, including 3 novel loci, 204 novel candidate genes and 69 replicated from GWAS. The 3 novel loci located at chr6: 72417543, chr15: 69601206, chr21: 30530692, mapping to gene *RIMS1*, *SPESP1*, *MAP3K7CL*. The results of colocalization analysis indicated that 142 of them showing strong evidence of a single shared causal variant and 134 of them showing evidence of joint causal variants. Their biological function was directly or indirectly associated with the occurrence of OP validated by VarElect tool. Several important OP-associated pathways were detected by protein-protein interaction and pathway enrichment analysis. Target genes were further enriched for differential expression genes in osteoblasts expression profiles, e.g. *IBSP*, affecting calcium and hydroxyapatite binding, and *CD44*, regulating alternative splicing of gene transcription. Transcriptome fine-mapping identifies more disease-related genes and provide additional insight into the development of novel targeted therapeutics to treat OP.

## Background

Osteoporosis(OP) is a highly polygenetic disease which has been studied intensively on the genetic level, resulting in abundant detections associated with gene loci and polymorphisms (Peacock et al. 2002; Clark and Duncan 2015). Osteoporosis is defined clinically that bone mineral density is 2.5 standard deviations or more below the young adult mean and remains the single golden standard predictor of primary osteoporotic fractures (Nguyen et al. 2007; Duncan and Brown 2010; Rachner et al. 2011). Bone mineral density is highly heritable, with evidence showing that the heritability of bone mineral density ranges from 50% to 80% (Duncan and Brown 2010). The recent large genome-wide association study (GWAS) to date estimated bone mineral density at the heel in 426,824 individuals and identified 1,103 independent genome-wide significant associations at 518 loci (Morris et al. 2019). They explain about 20% phenotypic variance in estimated bone mineral density. However, the majority of GWAS hits are in non-coding regions and their biological methnasims are difficult to understand (Moonesinghe et al. 2008; Nicolae et al. 2010).

The effect of genetic variation on phenotype is complex, where it may alter the abundance of one or more proteins by regulating gene expression and then affects the trait (SNP-Expression-Phenotype) (Musunuru et al. 2010; Lappalainen et al. 2013; Westra et al. 2013; Albert and Kruglyak 2015; Zhang et al. 2015). Gene expression is arguably the most impactful and well-studied effect of regulatory genetic variation. GWAS loci are enriched for expression quantitative trait loci (eQTL), rendering it a potential link between genetic variant and biology of disease (Stranger et al. 2007; Gusev et al. 2014; Lee et al. 2015). While most GWAS studies do not concomitantly measure gene expression, the influence of genetic variation on gene expression allows us to use gene expression reference datasets to predict gene expression given a set of genotypes, and subsequently identify new disease-associated genes (Nica et al. 2010; Nicolae et al. 2010; Albert and Kruglyak 2015). Transcriptome-wide association study (TWAS) approach has been implemented to identify genes with expression associated with complex traits by integrating genetic and transcriptional variation (Gusev et al. 2016; Barbeira et al. 2018). Instead of testing millions of SNPs in GWAS, TWAS evaluates the association of predicted expression for thousands of genes, greatly reducing the burden of multiple comparisons in statistical inference. This approach has been shown to have the potential to identify the genes responsible for GWAS-identified associations for complex traits and provide mechanistic insight regarding genes being regulated via disease-associated genetic variants (Mancuso et al. 2017; Gusev et al. 2018; Lu et al. 2018; Wu et al. 2018; Atkins et al. 2019).

In this paper, we conducted transcriptome-wide association study to identify genes associated with OP by integrating gene expression from the Genotype-Tissue Expression (GTEx) and GWAS summary data from the Genetic Factors for Osteoporosis (GEFOS) Consortium, and then evaluated the biological patterns of expression-trait association by COLOC method. Next, we performed VarElect to understand the biological function of association between the TWAS-significant genes and OP. Comparing with the results of differential analysis of the two mRNA expression profiles for OP, we further verified the causal associations between OP and TWAS-significant genes.

## Results

### TWAS identified candidate genes for OP

We performed TWAS method. Two gene-expression reference panels in muscle-skeletal and whole blood were used, with totally 13,416 genes towards GWAS summary data from GEFOS consortium to identify novel genes associated with OP. TWAS identified 276 significant associated genes at p-value < 3.7E-6, as shown in Figure 1.

**Figure 1.**
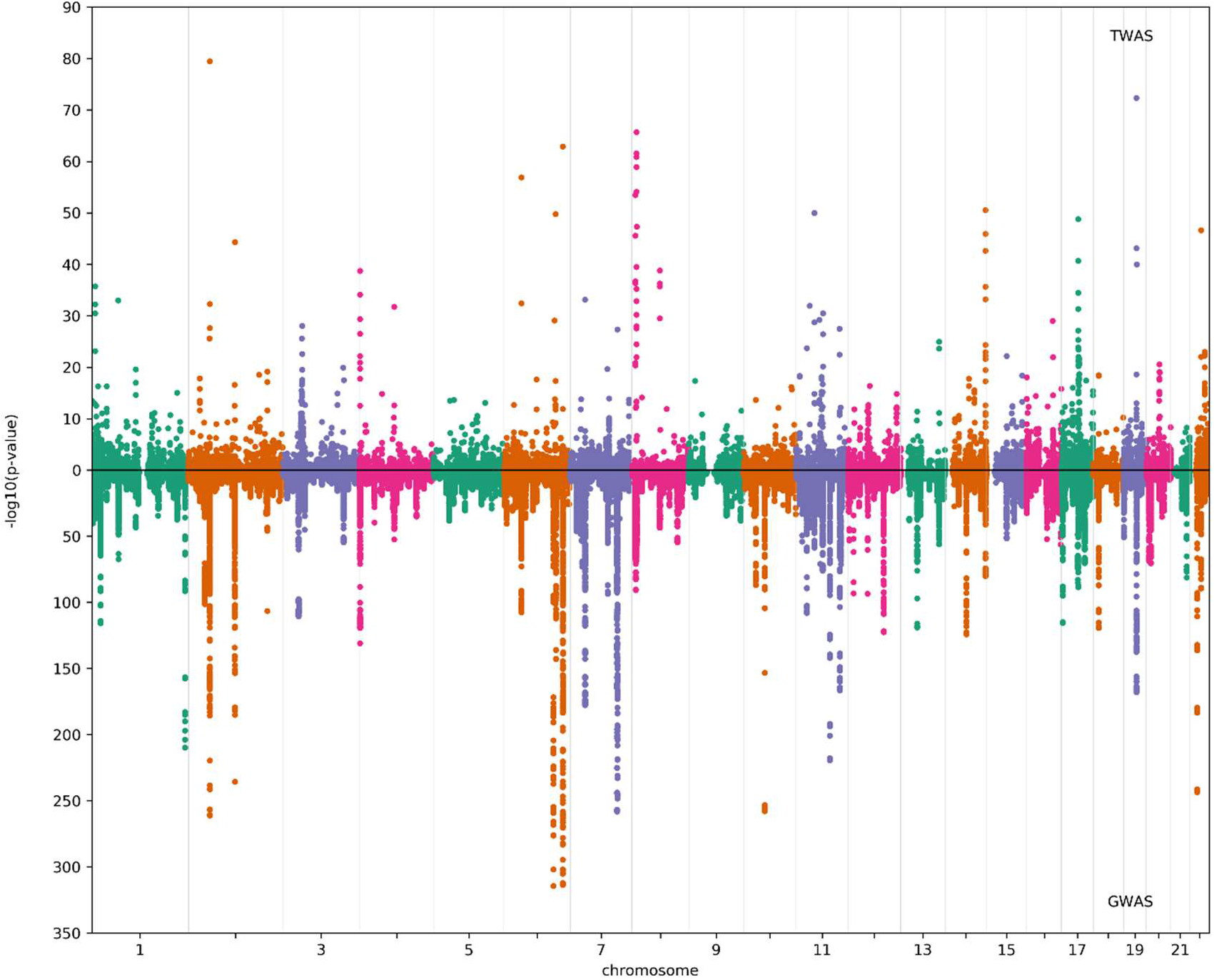
A manhattan plot of the results from TWAS analysis and GWAS analysis for OP. The transcriptome–wide significance threshold is p-value = 3.7E-6; The genome–wide significance threshold is p-value = 6.6E-9. There are 1,103 conditionally independent SNPs at 515 loci passed the criteria for genome-wide significance in n = 426,824 UK Biobank participants.

TWAS method can detect causal genes by introducing the effect prediction of genetic variants on the gene expression. There are the following four biological patterns identified by TWAS (Figure 2). First, for significantly associated SNPs with OP in the coding regions (introns and extons), the causal genes identified by GWAS and TWAS are more likely to be consistent, as shown in Figure 2a. The effect size of rs10411210 (P_GWAS_ = 1.6E-119) on OP in GWAS is corresponding with that of rs10411210 on *RHPN2* (P_TWAS_ = 4.4E-73) gene expression in TWAS. Second, for SNPs in non-coding regions, the candidate genes may be close to the significant eQTLs but different from the GWAS hits, as shown in Figure 2b. Variant rs2785197 (P_GWAS_ = 6.5E-44) in 11p13 mapping to *PDHX* in GWAS, but the causal gene for rs2785197 is more likely to be *CD44* (P_TWAS_ = 1.1E-32) in our TWAS. The colocalization analysis showed *CD44* (PP4=0.99 in Supplementary Table 3) gene expression was regulated by single variant rs2785197, which may be regarded as its expression regulation element. Third, the candidate genes may be regulated by relatively distant significant SNP in non-coding regions, as shown in 2c. Our TWAS results indicated that rs4792909 (P_GWAS_ = 1.5E-74) in 17q21.31 may be associated with *G6PC3* (P_TWAS_ = 4.2E-26). The distance between rs12478002 and *G6PC3* was 349kb, but we did not found gene reported by GWAS near rs4792909. Fourth, the candidate genes were discovered in non-significantly associated SNPs with OP. There GWAS non-significants regions as novel loci: rs1003260 (P_GWAS_ = 3.6E-08) in the 6q13 associated with *RIMS1* (P_TWAS_ = 2.1E-8) shown in Figure 2d, rs12917011 (P_GWAS_ = 2.1E-06) in the 15q23 associated with *SPESP1* (P_TWAS_ =3.3E-8) shown in Supplementary Figure 6a, rs2251381 (P_GWAS_ = 1.4E-06) in the 21q21.3 associated with *MAP3K7CL* (P_TWAS_ = 1.1E-9)shown in Supplementary Figure 6b. These 3 novel discoveries are firstly reported to be associated with BMD and further investigation can be performed.

**Figure 2.**
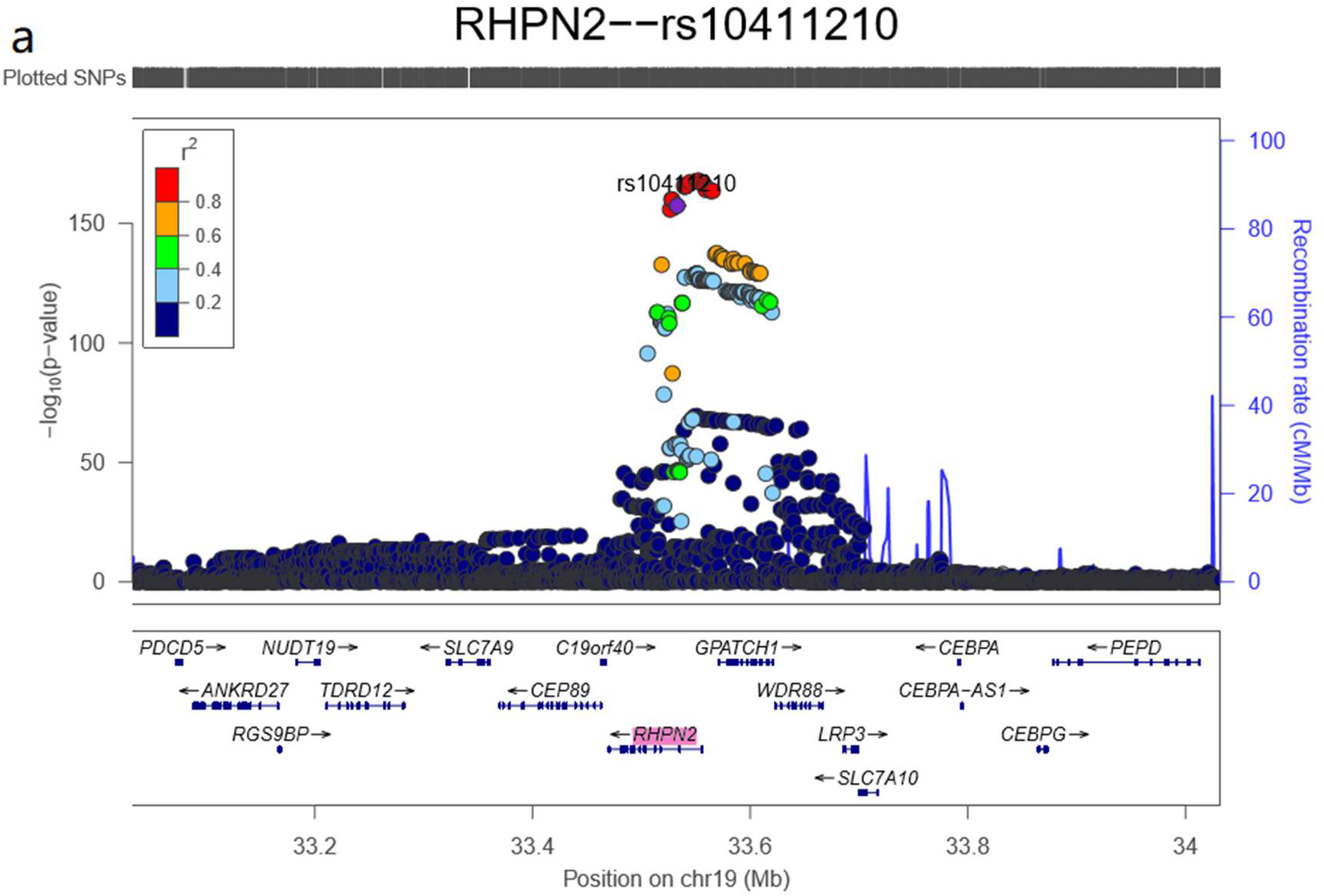
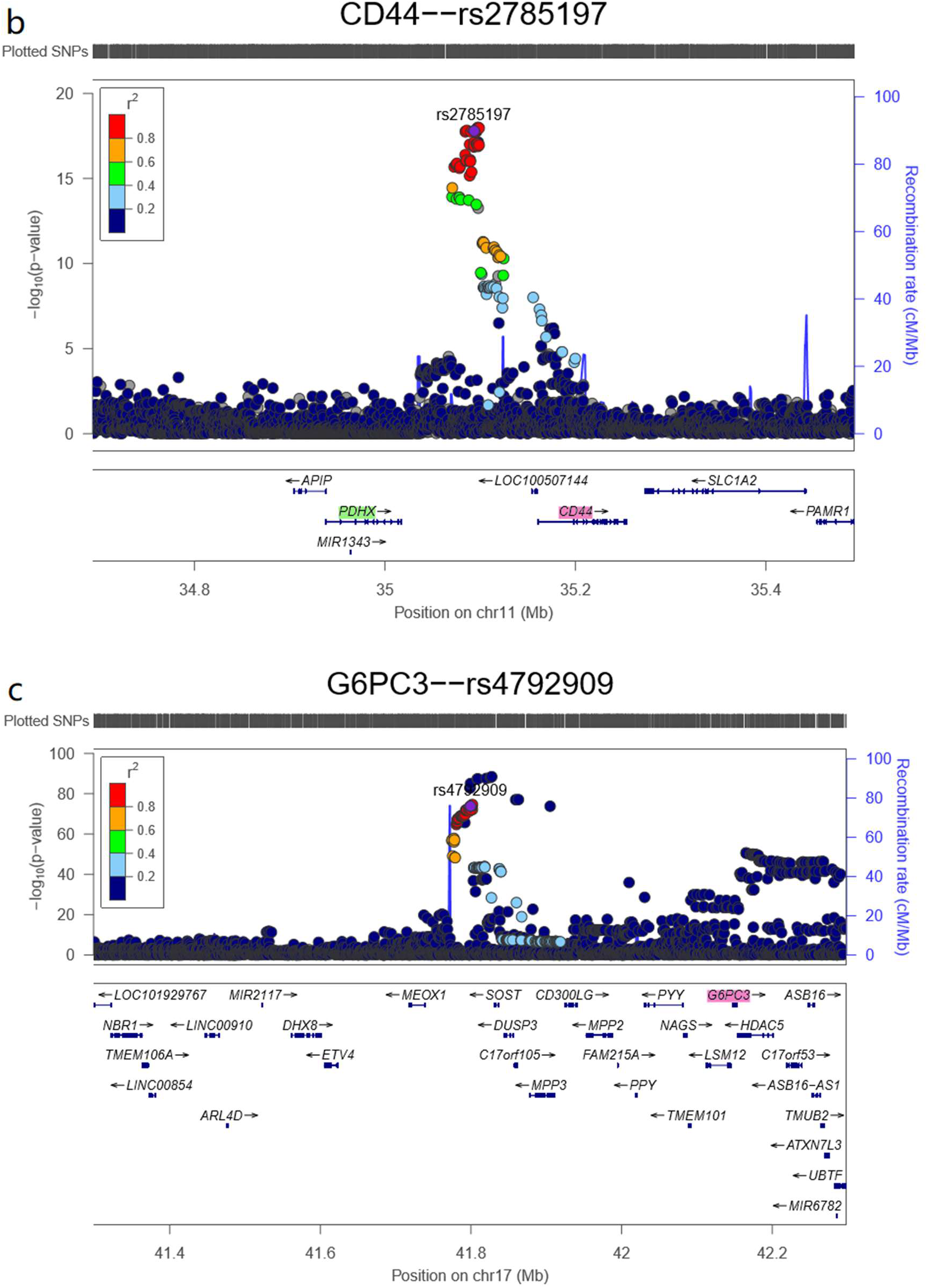
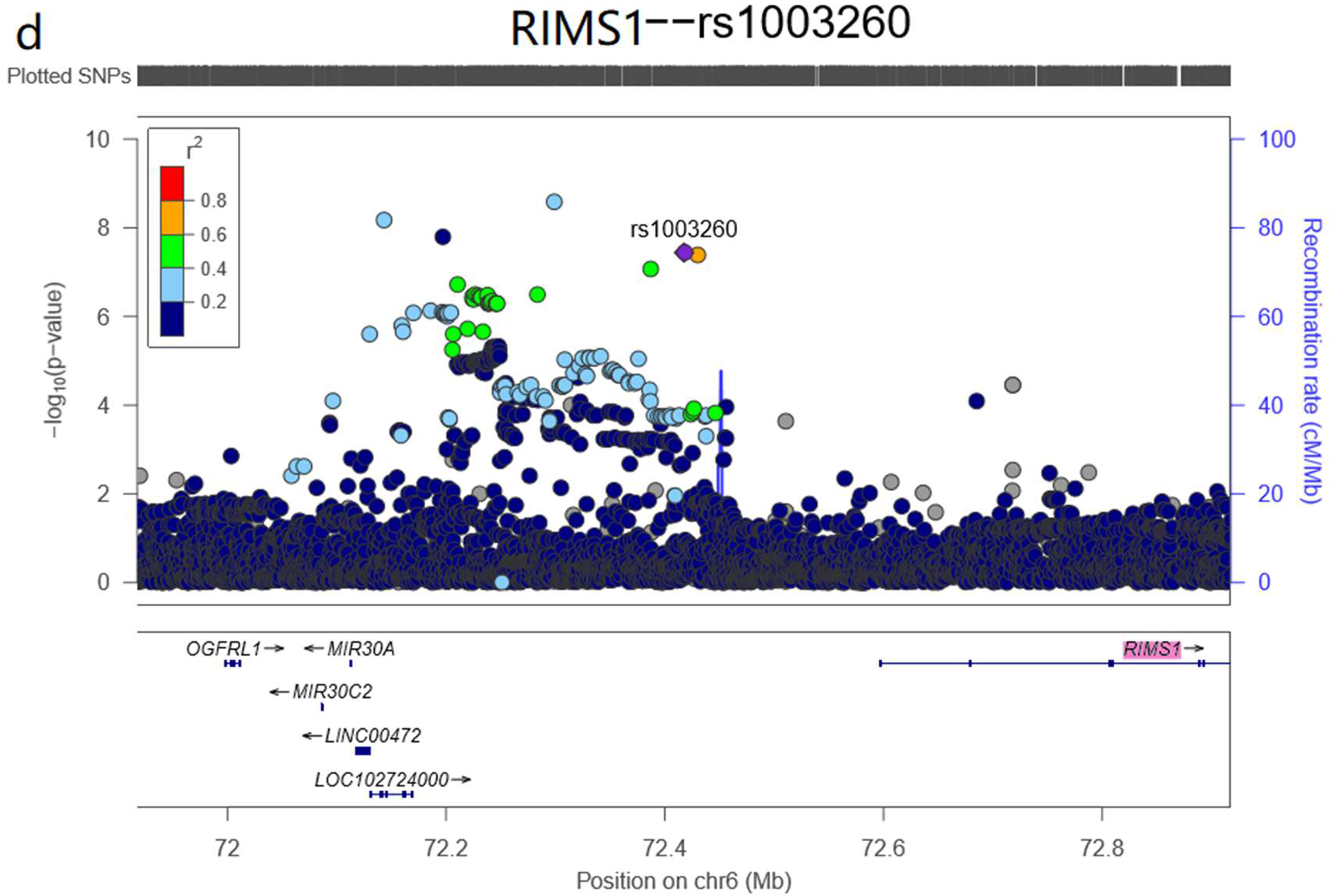
Biological patterns identified by TWAS. a: For significant SNPs in the coding regions, rs10411210 (P_GWAS_ = 1.6E-119) in 19q13.11 is associated with *RHPN2* (P_TWAS_ = 4.4E-73). b: For SNPs in the non-coding regions, rs2785197 (P_GWAS_ =6.5E-44) in 11p13 associated with *PDHX* marked green in GWAS, but The causal gene for rs2785197 is more likely to be *CD44* marked red (P_TWAS_ = 1.1E-32) in our TWAS. c: rs4792909 (P_GWAS_ = 1.5E-74) in 17q21.31 may be associated with *G6PC3* (P_TWAS_ = 4.2E-26). The distance between rs4792909 and *G6PC3* was 387kb, but there is not found gene reported by GWAS near rs4792909. d: rs1003260 (P_GWAS_ = 3.6E-08) in the 6q13 associated with *RIMS1* (P_TWAS_ = 2.1E-8). rs1003260 is not significant in GWAS.

Genes expression difference identified by TWAS may be causally associated with the phenotype of interest, but also can be due to variants LD or co-expressions (Huang et al. 2015; Hormozdiari et al. 2016). To pinpoint causal relationship between the target gene of an eQTL and the complex trait, we performed colocalization analysis by using COLOC method; see Methods section. The results showed that 134 TWAS associations have strong evidence of joint causal variants with PP3 > 0.9 shown in Figure 3a and Supplementary Figure 1 and Table 1, and 142 have evidence of a single shared causal variant with PP4 > 0.8 shown in Figure 3a and Supplementary Figure 1 and Table 2.

**Figure 3.**
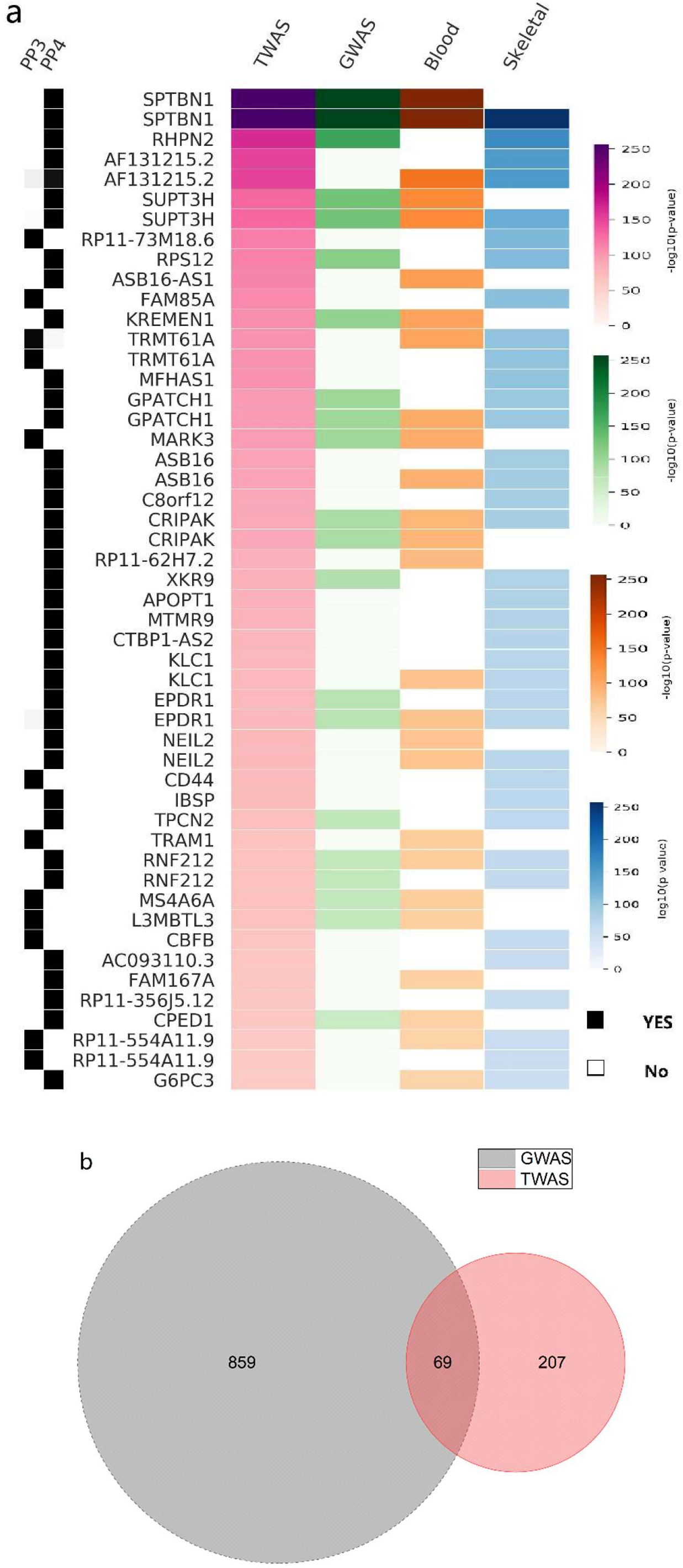
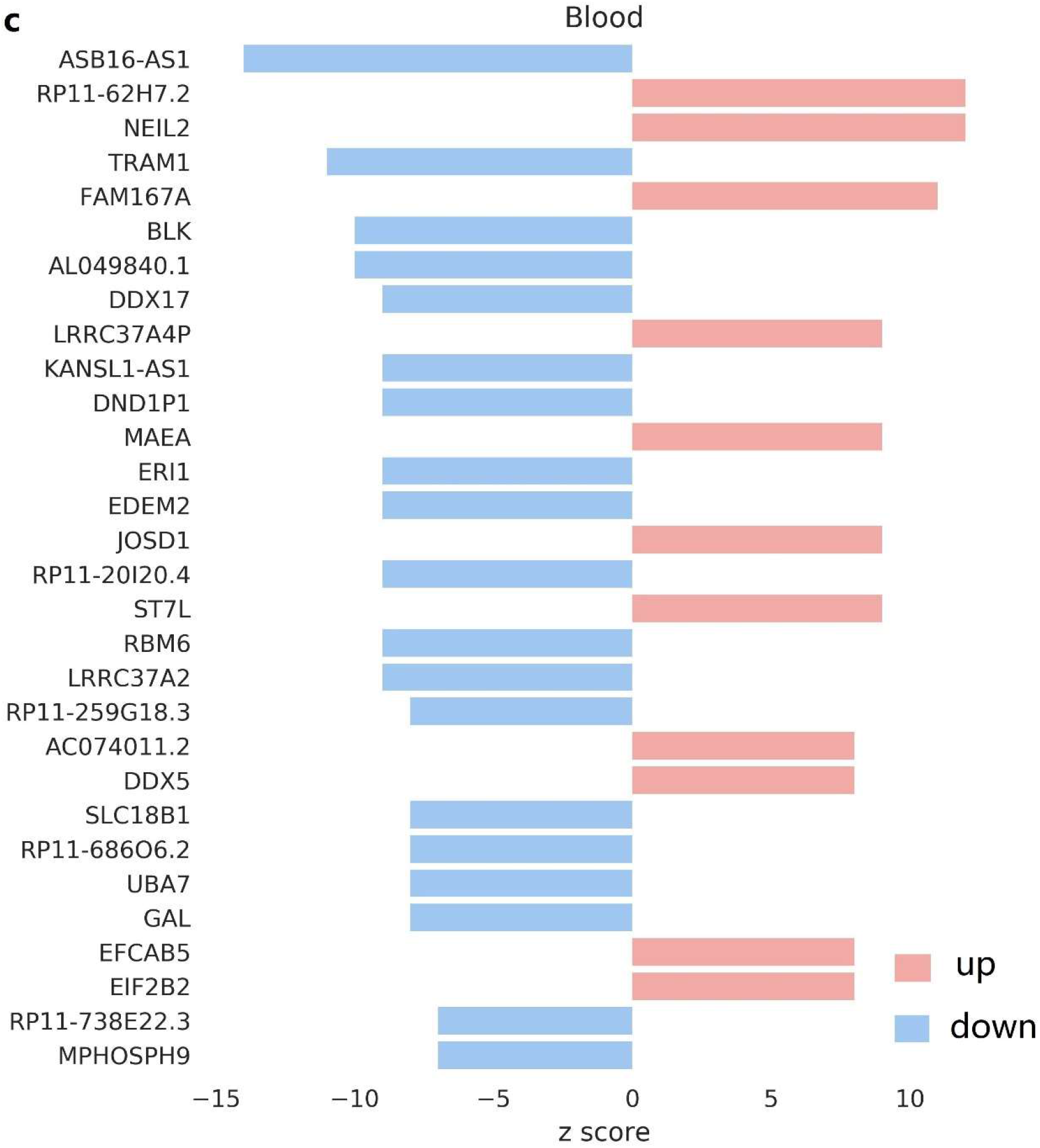
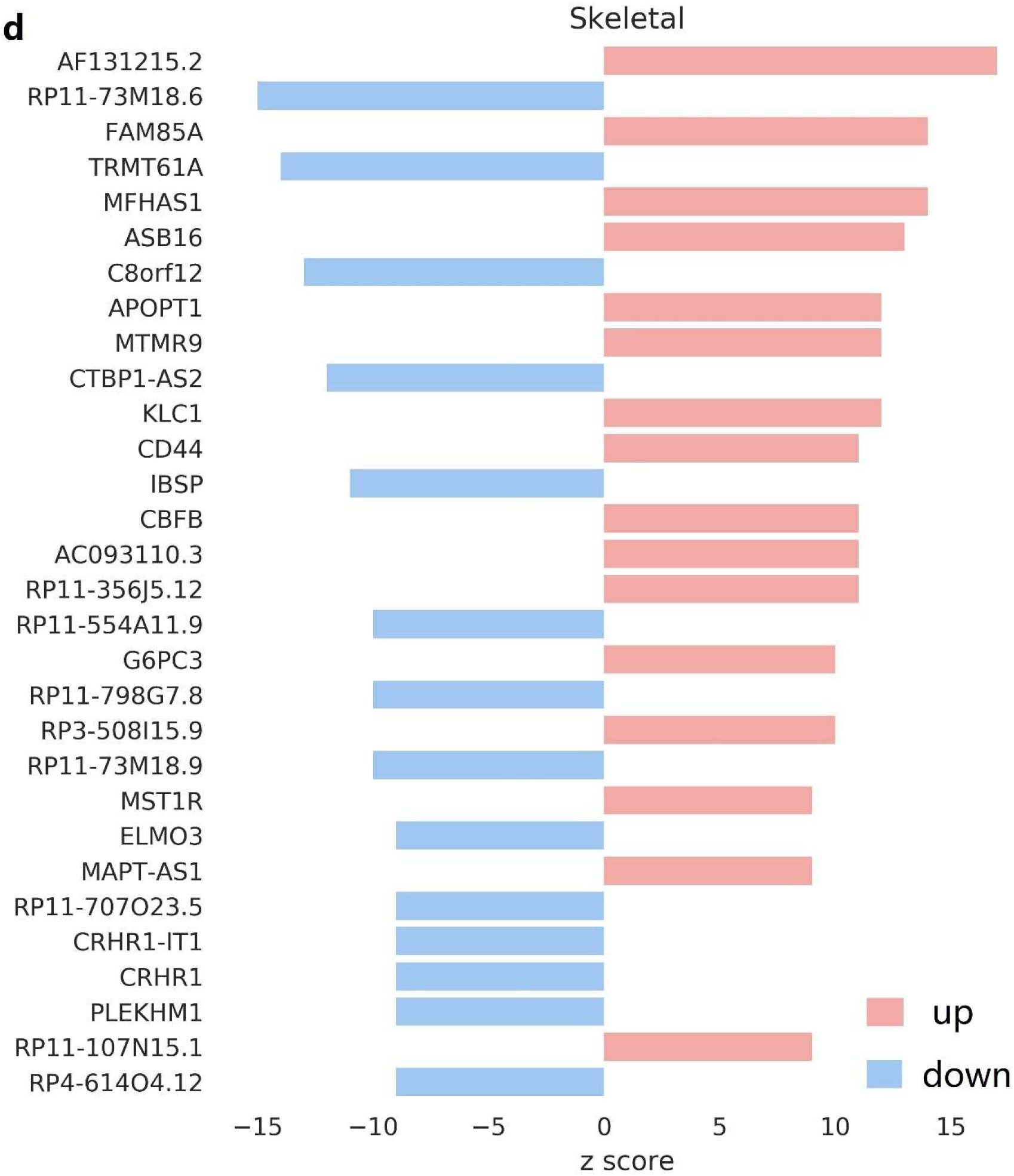
TWAS–significant genes and novel candidate genes in blood and skeletal tissue. PP3: represents trait–gene expression associations are caused by two distinct causal variants. PP4: represents trait–gene expression associations are caused by a signale causal variants. a: Heatmap of top 50 TWAS-significant genes, whether discovered in previous GWAS study, blood tissue or skeletal tissue, whether passed the criteria for colocalization analysis in PP3 > 0.9 or PP4 > 0.8 (full lists can be found in Supplementary Figure 1, Supplementary Table 2 and 3). b: Comparison of associated genes found by TWAS and GWAS methods. c and d: Top 20 novel candidate genes were found in blood and skeletal tissues respectively, the red bars represent gene expression up-regulated, and blue bars indicate down-regulated (full lists can be found in Supplementary Figure 2 and 3, Supplementary Table 2 and 3).

Comparing with previous GWAS studies, we observed that 3 genes located in novel loci and 204 genes have not been reported to be associated with OP risk in previous GWAS loci, 69 genes were previously implicated to be OP risky by literature using either GWAS or functional studies shown in Figure 3a-3b. We also found that 117 (117/276) genes were expressed in skeletal tissue and 71 (71/276) genes were expressed in blood tissue, and 88 (88/276) genes were expressed in both tissues as shown in Figure 3a. For 69 genes found in previous GWAS studies, our results provided additional evidence to support these previous findings. For the rest genes, 79 in blood tissue and 128 in skeletal tissue were considered as novel candidates shown in Figure 3c-3d.

### Assessment of the candidate gene-OP association

For 207 novel candidate genes, we evaluated the associations between the candidate genes and OP by implementing VarElect analysis. The analytical results showed that 24 genes (Supplementary Table 3) were ‘direct’ associations and 129 genes were ‘indirect’ associations (Supplementary Table 4); the rest were unclassified yet. The direct associations indicated the target genes were supported by rich evidences (the relevant literature, gene function annotation, etc.). The score in Supplementary Table 3 indicated the strength of the association between the gene and OP: the higher score, the stronger evidence. Indirectly associated genes may interact with intermediary to influence the development of OP, though protein interaction netorks and pathways (Supplementary Table 5). The remaining uncharacterized genes are mainly lncRNA, transcripts as the potential disease factors without available evidence requiring further investigations.

### Functional pathways of the candidate genes

In order to further verify the associations between the TWAS-significant genes and OP, we explored the biological function pathways of these genes by applying STRING and CluePedia tool. We found the majority of pathways were easy understanding to the occurrence of OP and some of them interact with each other (e.g. focal adhesion and ECM-receptor interaction, PI3K-Akt signaling), as shown in Figure 3a and Supplementary Table 5. We further enriched the functional pathways for the categorized gene lists (direct, indirect, in blood, in skeletal). We found all significant pathways (p-value < 0.5) were enriched in the skeletal tissue while part enriched in blood. ‘Direct’ genes can be enriched in the critical pathway such as mineral absorption and calcium signaling pathway. These results showed that TWAS–significant genes involved many biological mechanisms in developing OP.

### Functional validation for the candidate genes

Previous research, utilizing expression profiling with gene signatures of cellular models to characterize the gene’s involvement in bone metabolism and disease processes, found that impaired osteoblastic differentiation reduces bone formation and causes severe OP in animals (Stein et al. 1990; Wu et al. 2003; Misof et al. 2012). We analyzed two gene expression profiles GSE35956 and GSE35959 from GEO, containing two groups people: the primary OP and normal. Based on the cut-off criteria of p < 0.05 and logFC > 1 to select DEGs, a total of 156 and 265 DEGs were identified from GSE35956 and GSE35959 datasets. Comparing DEGs with TWAS-significant genes, 5 up-regulated and 2 down-regulated genes overlapped in two type datasets shown in Supplementary Figure 4. We observed that these genes were quite significant in TWAS, and their expression differences were also consistent with COLOC analysis results, as shown in Figure 4a-4d. Therefore, we inferred that these genes are very likely to be the causal pathogenic gene of OP. The results of functional pathway analysis also supported our findings, as shown in Figure 4e-4g, *IBSP* and *CD44* included in the ECM-receptor interaction pathway, which is a branch of the focal adhesion pathway and acts on PI3K-AKT signaling pathway. Owning to the small samples size of gene expression datasets, more experiments are needed in the future.

**Figure 4.**
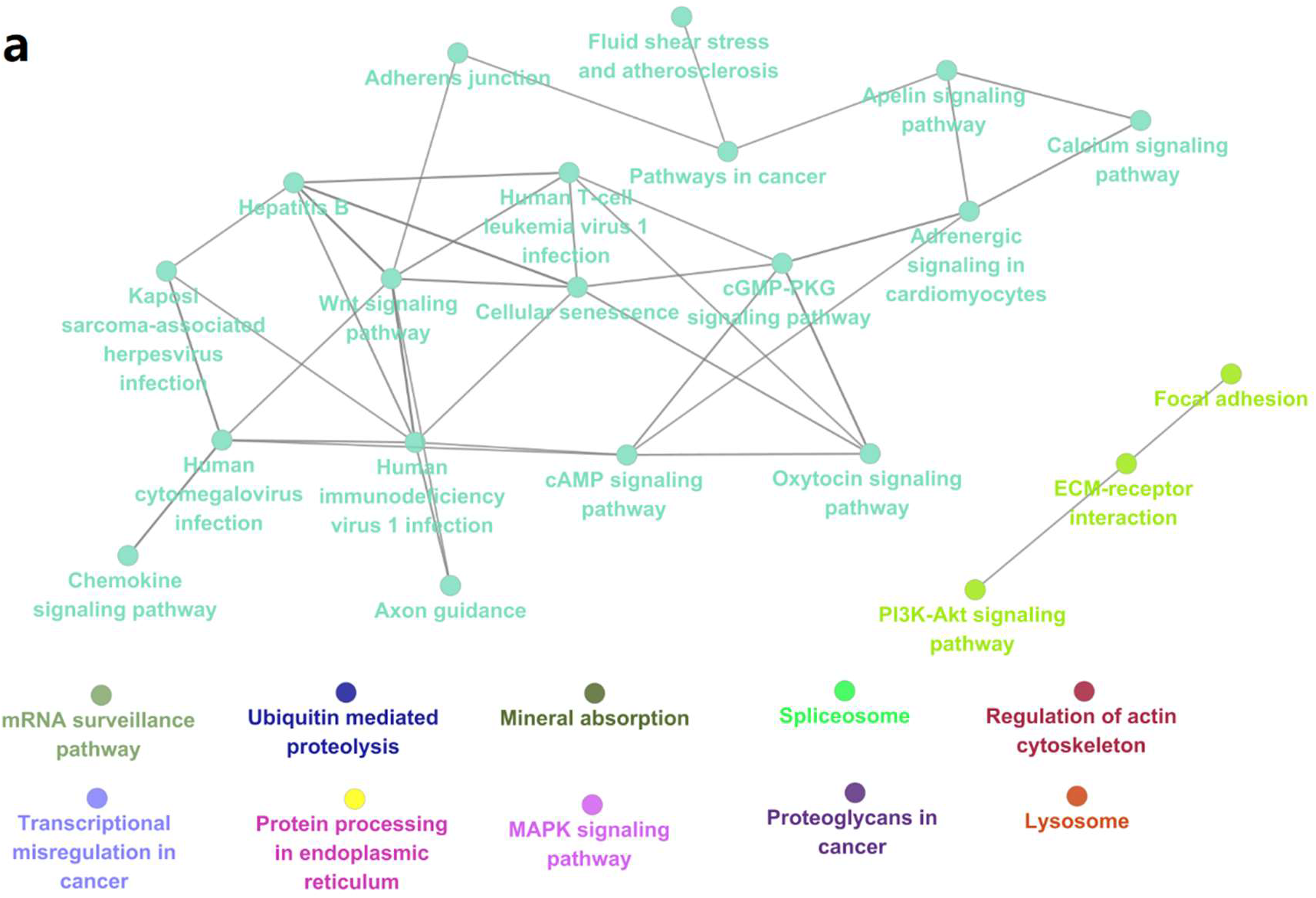
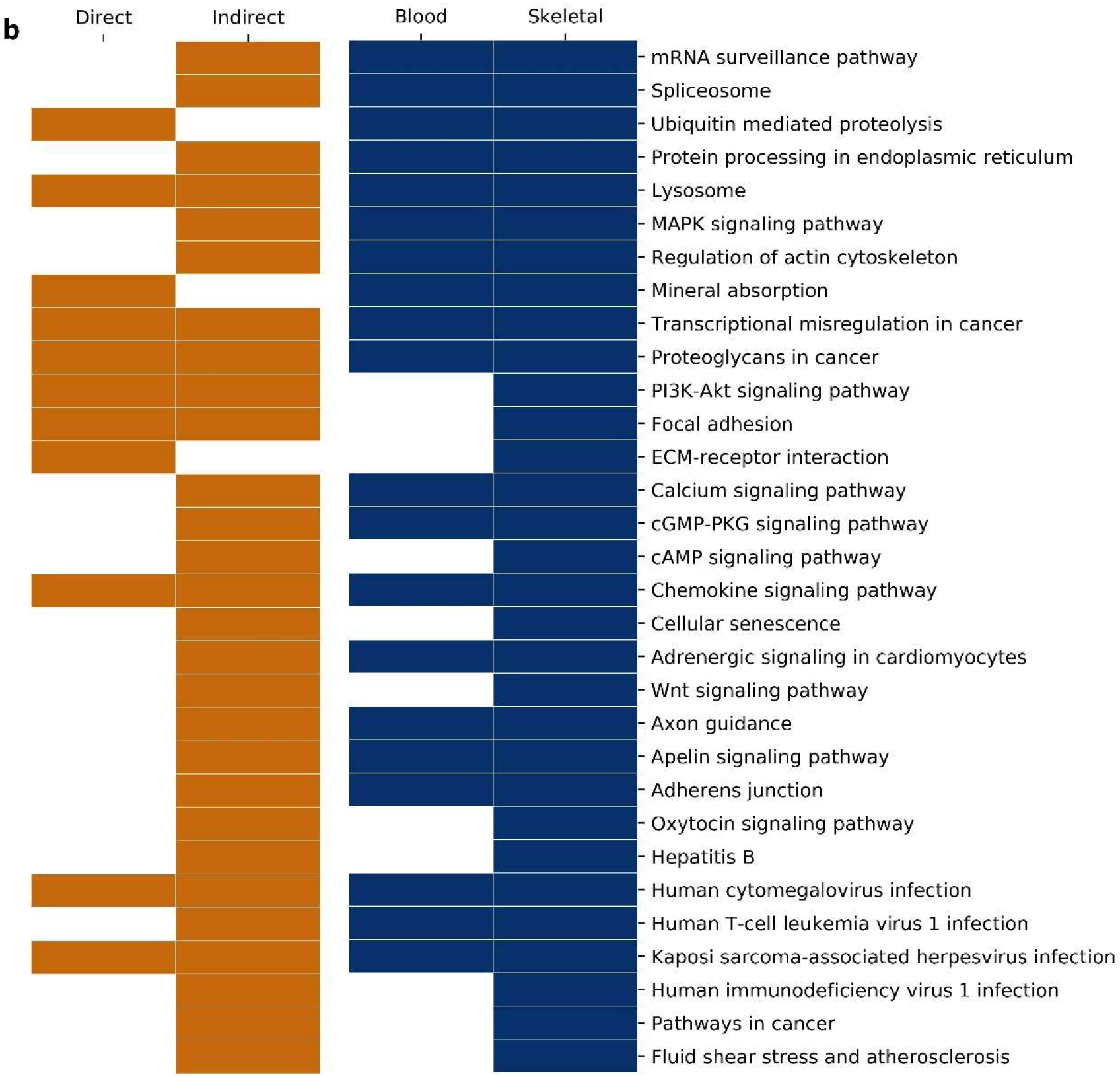
The functional pathways of TWAS-significant genes. They are enriched by applying STRING and CluePedia tool. Significance of pathways is determined by the hypergeometric test (one-sided) followed by Fisher’s combined probability test (one–sided) to determine combined pathway significance (p–value < 0.5). a: The functional pathways of TWAS–significant genes, the circles represent functional pathways, and the line represents the interactions between pathways. b: Classification of functional pathways according to the categorized gene lists. Genes in skeletal tissue are enriched in all significant pathways. The genes list for each pathway are found in Supplementary Table 5.

**Figure 5.**
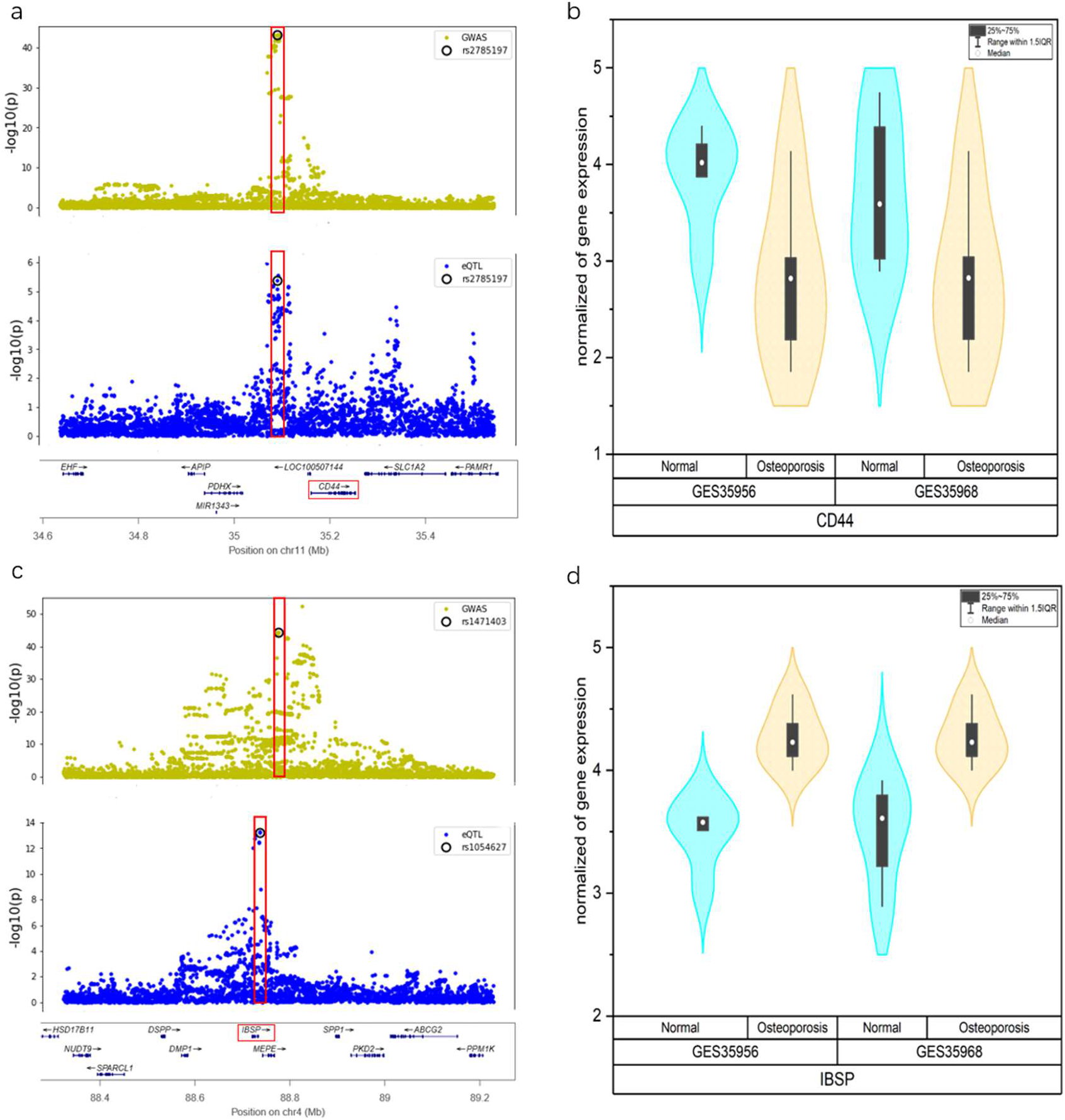
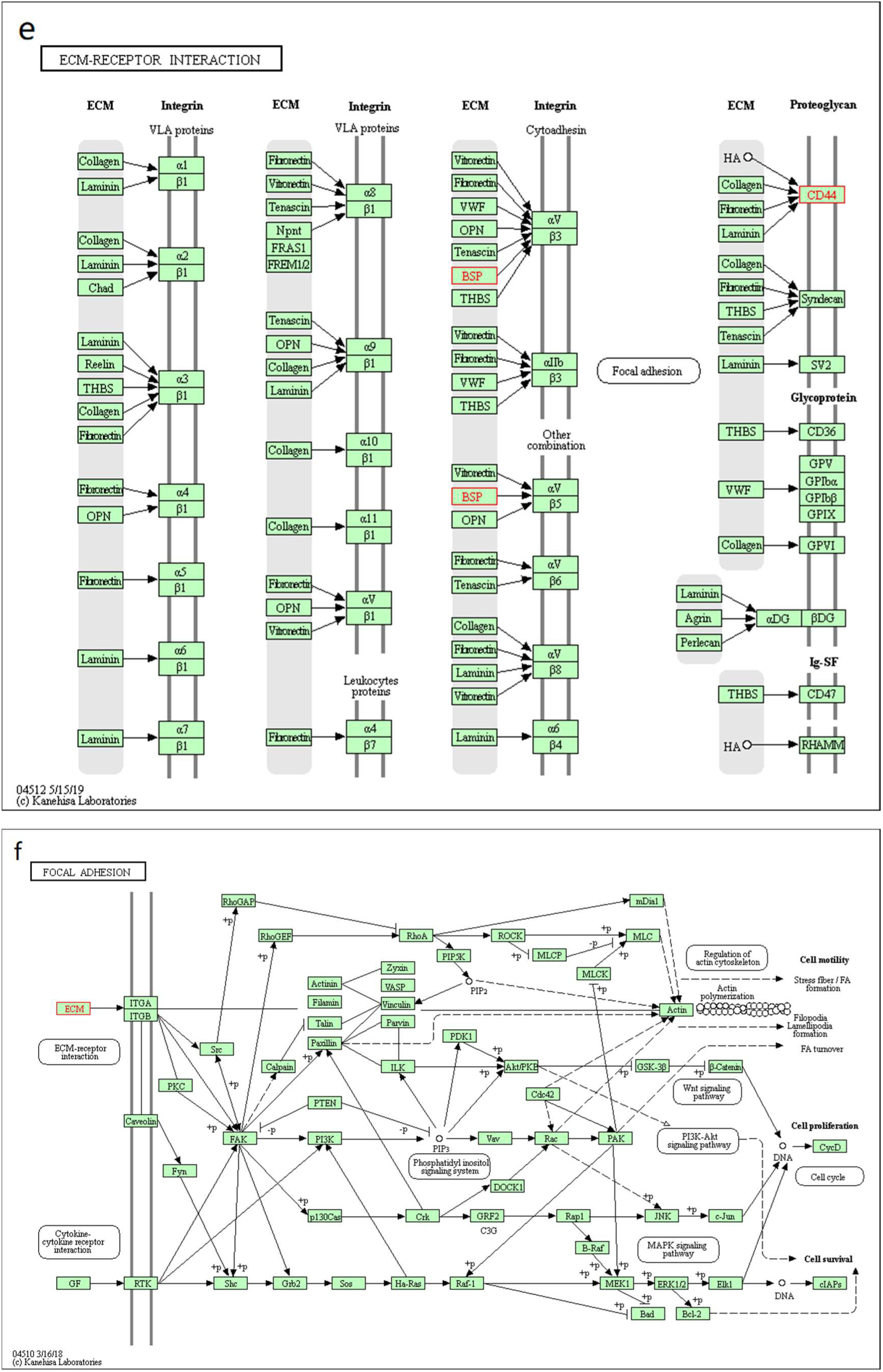
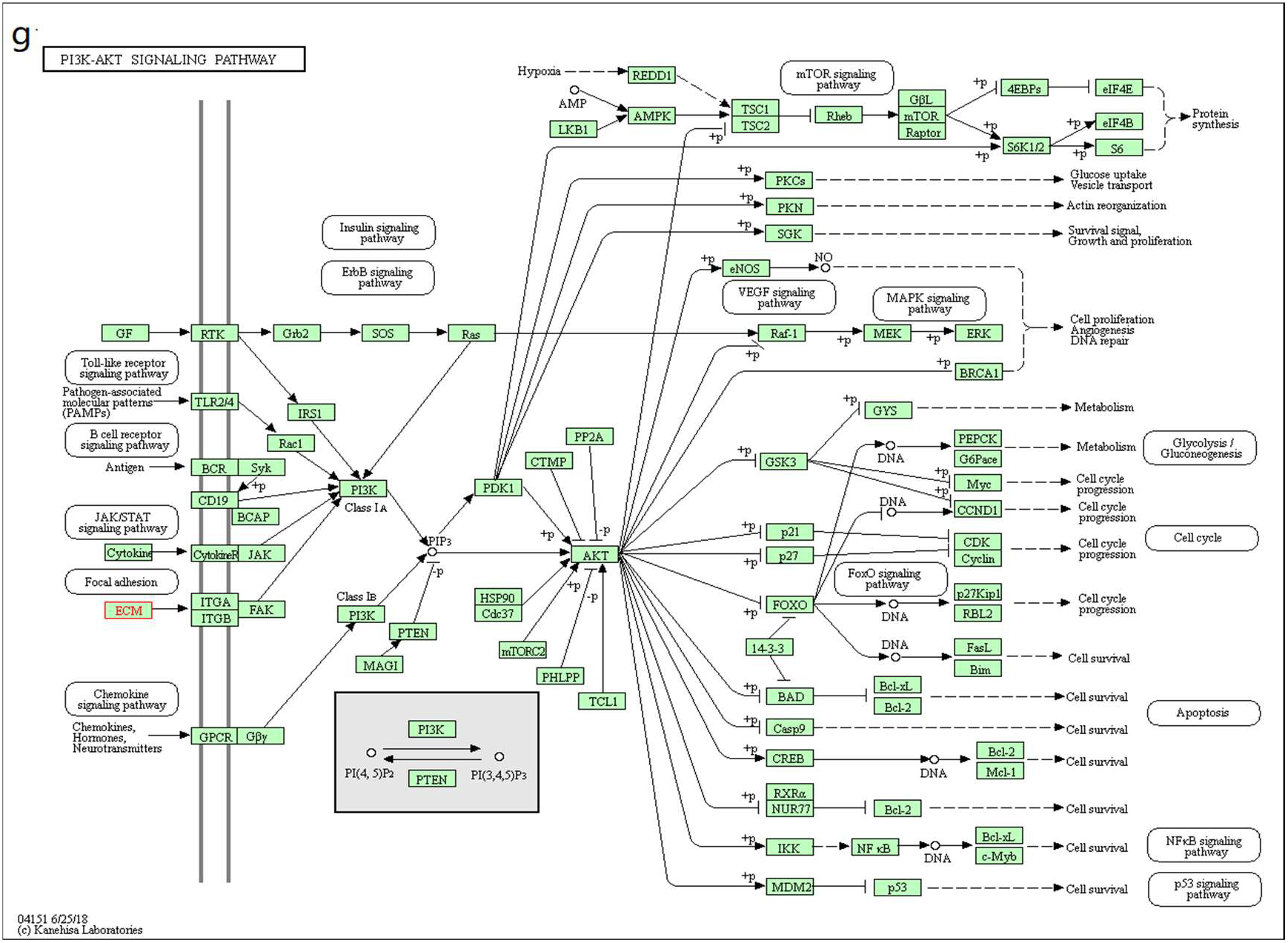
Biological function verification for genes. TWAS–significant gene *CD44* (P_TWAS_ = 1.1E-32) and IBSP (P_TWAS_ =1.8E-32) are validated by COLOC method, gene expression profiles, and biological function pathways. a and c: The colocalization analysis results for gene *CD44* and *IBSP*, showing a single shared causal variant rs2785197 and joint causal variants rs1471403 and rs1054627 respecively. b and d: Gene expression for *CD44* and *IBSP* in the GSE35956 and GSE35959 datasets. e-g: *IBSP* and *CD44* are enriched in the ECM-receptor interaction pathway, which is a branch of the focal adhesion pathway and acts on PI3K-AKT signaling pathway, also see Figure 4a.

### TWAS for OP identifies new loci

We found 3 genes in novel loci. *RIMS1* (P_TWAS_ = 2.1E-8) associated with rs1003260 (P_GWAS_ = 1.8E-8, MAF = 0.125) and is a *RAS* gene superfamily member in 6q13 that regulates synaptic vesicle exocytosis; *SPESP1* (P_TWAS_ =3.3E-8) in 15q23 associated with rs12917011 (P_GWAS_ = 2.1E-06, MAF = 0.438) code a human alloantigen involved in spermegg binding and fusion; *MAP3K7CL*(P_TWAS_ = 1.1E-9) associated with rs2251381 (P_GWAS_ = 1.4E-06, MAF = 0.367) and is a protein coding gene in 21q21.3. VarElect analysis showed that the biological function of three genes were indirectly associated with OP and provided evidence for causality for OP shown in Supplementary Table 4. However, we did not enrich significant functional pathways for three genes, the causal effect of them on OP needs to be verified by advanced biological experiments. As shown in Figure 2d and Supplementary Figure 6a-6b, we observed that the distance between the three causal SNPs and causal genes is within 500kb, and other significant GWAS SNPs were not found, the results indicated one of the advantage of TWAS method, which can find causal genes in the non-significant GWAS regions.

## Discussion

Multiple GWAS studies have been performed with considerable sample sizes to detect OP heredity, yet progress towards understanding disease mechanisms has been limited. Most GWAS hits are in non-coding regions and difficult to understand the downstream biological inference. In most cases, the nearest genes were uaually reported (Smemo et al. 2014; Claussnitzer et al. 2015; Spain and Barrett 2015). In fact, SNPs in the noncoding region did not have to regulate gene based on the distance between SNPs and genes. Intergreting GWAS data and transcriptome data will empower novel discovery and possibly pinpoint the causality. TWAS method calculated local SNPs-gene expression correlations, and further calculated likelihood of genes causality. Therefore, for a significant SNPs in the coding region, the causal genes identified by GWAS and TWAS should be and indeed are consistent, as shown in Figure 2a. For SNPs in noncoding regions, the causal genes may be close to the significant eQTLs but different from the GWAS hits as shown in Figure 2b. TWAS method can even discover causal genes non-significantly associated SNPs with OP shown in Figure 2c, and relatively distant significant SNP shown in Figure 2d. More valuable region plots can be found in Supplementary Figure 5-6.

We totally found 276 candidate genes, of which 69 were replicated in GWAS, and the rest 207 were novel candidates. Among them, 142 target genes are regulated by two distinct causal variants, and 134 target genes share one causal variant. By analyzing the biological functions behind, we found that 24 novel candidate genes directly affect the pathways closely related to the development of osteoporosis in our results: *IBSP, EIF2B2, CD44*, *FEN1, UBA7*, *MARCO*, *ATF1, CBFB*, *G6PC3*, *SLC11A2*, *GAL*, *CCR3, MST1R, PLEKHM1*, *ATRIP*, *CCDC36*, *AKAP7*, *EPRS*, *CTSB*, *ASB16-AS1*, *CRHR1*, *FADS1*, *MAP1LC3A*, *MAEA*. For example, *SLC11A2* enriched in mineral absorption pathway regulates the fine-tuned balance between bone resorption and bone formation and thus affects bone density (Xu et al. 2017), shown in Supplementary Figure 7. In the other hand, 129 novel candidate genes seem exerting their biological functions to affect the development of osteoporosis through protein-protein interaction networks. As shown in Supplementary Figure 8, *RAC3* and *NFATC4* were enriched in the MAPK signaling pathway through interacting with genes (*ESR1*, *FOS*, *IGF1*, *TGFB1*, *JUN, NFATC1*, *IGF*1, *LRP5*, *TNF*, *PRKACA*) known to be associated with osteoporosis. MAPK signaling pathway is involved in the regulation of many cellular physiological functions such as proliferation, differentiation, inflammation, and apoptosis, and affect bone formation (Peng et al. 2009; Wanachewin et al. 2012). More information on gene interactions can be found in Supplementary Table 5.

We found *RIMS1*, *MAP3K7CL*, *SPESP1* located in new loci and their causal SNPs were non-significantly associated with OP in GWAS. *RIMS1*, regulating synaptic membrane exocytosis 1, is a RAS gene superfamily member and plays a role in the regulation of voltage-gated calcium channels during neurotransmitter and insulin release. *MAP3K7CL, MAP3K7* C-terminal like, is a protein coding gene. The GO annotation (GO:0005515) showed *MAP3K7CL* interact selectively and non-covalently with any protein or protein complex. But there is little research on its biological function. *SPESP1* code sperm equatorial segment protein 1 involved in fertilization ability of sperm. The current studies have not supported evidence for the causal association between three genes and OP, so we hope to have follow-up experiments to verify them.

Furthermore, we provided additional evidence by comparing with differential expression genes by analyzing two gene expression profiles in OP and non-OP groups. We found seven significant differential expression genes in our results: *IBSP, CD44, SPTBN1, PAPSS2, TRAM1, PPP1CB, NCKAP1*, shown in Supplementary Figure 4. *IBSP* is remarkably downregulated and associates with OP significantly (TWAS p-value=1.8E-32). SNP rs1471403 and rs1054627 may co-regulate gene expression of *IBSP* (PP3=1, Figure 4c-4d). Previous studies showed that IBSP is expressed in all major bone cells including osteoblasts, osteocytes and osteoclasts (Trošt et al. 2010) and encodes a major non-collagenous bone matrix protein binding to calcium and hydroxyapatite via its acidic amino acid clusters (Mafi Golchin et al. 2016). Another discovery *CD44* is remarkably upregulated. Previous research argued that a linkage synonymous mutation in exon 9 of the *CD44* gene through a cell experiment, may increase the susceptibility of the family to OP by influencing alternative splicing of gene transcription (Vidal et al. 2009). Information about other genes can be found in Supplementary Table 6.

This is as yet largest study integrating GWAS and TWAS to identify susceptibility genes of OP. We used data from the 426,824 individuals GWAS of OP and 860 samples TWAS in our analyses. Many findings were discovered, although there still exist limitations of this research. First, TWAS method cannot explain the variants influencing disease that are independent of cis expression, as it was only trained on cis-eQTL analysis. Second, there may be bias using normal blood and skeletal tissues from GTEx to make predictions. Third, tissue sensitivity and tissue specificity are important issues when running TWAS. Prediction models built on gene expression data from osteoblasts cells in OP patients will help identifying additional candidate genes associated with OP (Orlic et al. 2007).

In summary, we integrated data from GWAS and transcripome expression to identify 276 candidate genes associated with OP; 69 of them were replicated from GWAS, and 204 novel candidate genes in loci reported by GWAS and 3 novel candidate genes in new loci. We analyzed biological patterns of those loci and explained their pathway interactions. We hope that our findings will provide novel insights into the future pathogenetic studies of OP.

## Methods

### GWAS summary datasets of OP

The GWAS summary statistics for OP was derived from GEFOS Consortium website (URL) in December 2018. The phenotype feature of OP was measured by bone mineral density estimated from quantitative heel ultrasounds. The large scale GWAS analysis for OP were performed in a cohort of 426,824 participants (55% female) from UK Biobank (Morris et al. 2019). Briefly, GWAS analysis was performed based on the HRC imputation panel (hg19) including about 14,000,000 SNPs with MAF ≥ 0.05% and acceptable imputation quality (info score > 0.3). A detailed description of sample characteristics, experimental design, and statistical analysis can be found in the published study (Sudlow et al. 2015).

### Integration of GWAS and gene expression

To integrate GWAS results and gene expression, we used TWAS method. We included two relevant reference transcriptome datasets in our analysis: whole blood and muscle-skeletal from GTEx v7. TWAS method integrated information from expression reference panels (SNP-gene expression correlation), GWAS summary statistics (SNP-OP correlation), and linkage disequilibrium (LD) reference panels (SNP-SNP correlation) to assess the association between the cis-genetic component of expression and trait (expression-OP correlation) (Gusev et al. 2016; Gusev et al. 2018), In practice, the effect sizes of cis-SNP-expression in the 500kb loci region were estimated with a sparse mixed linear model (Zhou et al. 2013). TWAS used pre-computed gene expression weights combined with GWAS summary statistics to calculate the association effect for each gene to disease. In this study, the gene expression weights of whole blood and muscle-skeletal were derived from the FUSION website (URL). The genes with significant association signals were identified at p-value < 3.7E-6 after strict Bonferroni correcting.

### Evaluation of trait-gene expression associations

To evaluate the reliability of TWAS analysis results and understand the biological mechanisms of trait–gene expression associations, we performed COLOC method (Giambartolomei et al. 2014). COLOC method uses asymptotic Bayes factors with summary statistics and regional LD structure to estimate five posterior probabilities: no association with either GWAS or eQTL (PP0), association with GWAS only (PP1), association with eQTL only (PP2), association with GWAS and eQTL but two independent SNPs (PP3), and association with GWAS and eQTL having one shared SNP (PP4). For each of the GWAS hits, we defined a 500kb region at either side of the index variant and tested for colocalization within the entire cis-region of any overlapping eQTLs (transcription start and end position of an eQTL gene plus and minus 500kb, as defined by GTEx) in two human tissues from GTEx v7. A signal with PP3 > 0.9 was considered the evidence for trait - gene expression associations caused by two distinct causal variants from GWAS and eQTL. A signal with PP4 > 0.8 was considered the evidence for trait-gene expression associations caused by a joint signal from GWAS and eQTL.

### Assessment of gene-disease associations

To assess the likelihood of functional genes which are more likely to be causal, VarElect (Stelzer et al. 2016a; Stelzer et al. 2016b), a cutting-edge Variant Election application for disease/phenotype-dependent gene variant prioritization, were used to assess the associations of biological function between the candidate genes and OP. VarElect provides a robust algorithm for ranking genes within a short list, and pointing out their likelihood associated with disease, and produces a list of prioritized, scored, and contextually annotated genes and direct links to supporting evidence and additional information. VarElect utilizes the deep LifeMap Knowledgebase to infer the ‘direct’ or ‘indirect’ association of biological function between genes and phenotypes. ‘Direct’ association between genes and disease has been supported by many studies that genes can directly affect the development of disease. ‘Indirect’ association between genes and disease are based on shared pathways, protein-protein interaction networks, paralogy relations, domain-sharing, and mutual publications.

### PPI network and pathway enrichment analysis

The functional networks of TWAS-significant genes with OP were further validated by STRING and CluePedia tool. STRING (Search Tool for the Retrieval of Interacting Genes, URL) is an online tool designed to evaluate the protein-protein interaction (PPI) networks (Szklarczyk et al. 2015; Szklarczyk et al. 2017). The CluePedia is a plugin of Cytoscape software and search for potential genes associated with the certain signaling pathway by calculating linear and nonlinear statistical dependencies from experimental data (Shannon et al. 2003; Bindea et al. 2013). The PPI networks of TWAS-significant genes was constructed by STRING. The functioal pathways were detected and visualized by CluePedia. The pathways were identified at p-value < 0.5 (Bindea et al. 2013).

### Differential analysis of gene expression

To further validate the functional causslity of candidate genes, we compared the candidate genes with differential expression genes (DEGs) in osteoblasts for osteoporosis sufferer. The original datasets comparing the gene expression profiles between OP and normal controls were downloaded from NCBI GEO databases (URL). Two gene expression profiles GSE35956 and GSE35959 were based on GPL570 (Affymetrix Human Genome U133 Plus2.0 Array, Affymetrix, SantaClara, CA, U.S.A). We performed robust multi-array average approach (Hochreiter et al. 2006) for background correction and normalization. The original GEO data were then converted into expression measures. Limma package (Smyth 2005) was used for determining DEGs between OP samples and non-OP samples (p < 0.05 and log2FC >1 as the cutoffcriterion).

## Data access

GWAS summary data are available in the Genetic Factors for Osteoporosis (GEFOS) Consortium (http://www.gefos.org/); gene expression weights of whole blood and muscle-skeletal were derived from the FUSION website (https://gusevlab.org/projects/fusion/); Two gene expression profiles are available in NCBI GEO databases under der accession number GSE35956 and GSE35959.

## URL

GEFOS: http://www.gefos.org/

Fusion: https://gusevlab.org/projects/fusion/

GEO: https://www.ncbi.nlm.nih.gov/geo/

STRING: https://www.string-db.org/cgi/

## Author’s contributions

PY and MZ conceived the project and designed the experiments. MZ, HF, JJ, LY, WS and YL analyzed the data. PY and MZ wrote the manuscript. All authors read and approved the final manuscript

## Acknowledgements

We are thankful to our institutes who provided their expertise that greatly assisted this research work.

